# Protein aggregation drives cell aging in a size-specific manner in *Escherichia coli*

**DOI:** 10.1101/2024.10.11.617345

**Authors:** Linda Wagener, Arpita Nath, Murat Tuğrul, Abram Aertsen, Ulrich K. Steiner, Audrey M. Proenca

## Abstract

Aging, the decline in physiological function over time, is marked by the intracellular accumulation of damaged components. It can be attributed to a trade-off between the investment into organismal maintenance and the production of high-quality offspring, where the parent accumulates damage over time and retains it upon reproduction, while the offspring is rejuvenated. Asymmetric damage partitioning has been observed even in simple unicellular organisms, such as *Escherichia coli* bacteria, that retain aggregates of misfolded proteins during cell division. However, recent studies presented conflicting evidence on the effect of protein aggregates on fitness, ranging from detrimental effects on cell growth to enhanced stress survival. Here, we show that the decisive factor driving growth decline in *E. coli* is not the presence of a protein aggregate, but the proportion of the intracellular space occupied by it. By following single-cell *E. coli* lineages expressing fluorescently labeled DnaK chaperones, we quantified damage accumulation and partitioning across generations in microfluidic devices. Our results suggest that the aggregation of damaged proteins allows cells to keep damage separate from vital processes and compensate for the lost intracellular space by growing to larger sizes. This process results in morphologically asymmetric divisions, a finding that counters the long-assumed symmetry of *E. coli* cell division. In line with other recent evidence, our findings point to a more complex role of protein aggregation, with implications for our understanding of the cellular mechanisms underlying aging as well as its evolutionary origins.

## INTRODUCTION

Organisms age due to a decline in fitness over time, losing physiological integrity and becoming increasingly more vulnerable to death (*1, 2*). Aging can be attributed to a trade-off between reproduction and somatic repair, in which the maintenance of the germline comes at the cost of an increased damage accumulation by a “disposable” soma (*3*). Although unicellular organisms lack a differentiation between soma and germline, similar processes of asymmetric damage partitioning have led to the emergence of aging in yeast (*4*), algae (*5*), and bacteria (*6, 7*). These single-cell organisms can provide a system where the progression of aging hallmarks can be observed and quantified *in vivo* (*1*), thus offering a means to elucidate the origins of organismal aging at the cellular level.

Symmetrically dividing prokaryotes, such as the rod-shaped model bacterium *Escherichia coli*, were once considered not to exhibit the aging phenotype. However, despite splitting into two cells that are morphologically and genetically identical, *E. coli* displays a deterministic physiological asymmetry (*7*–*9*). Cells inheriting an old pole for multiple generations age, exhibiting slower growth rates, decreased offspring production, and increased mortality rates (*7, 10*). In contrast, cells inheriting a new pole, which was synthesized during the previous fission event, rejuvenate. Because this asymmetry presumably arises through the uneven partitioning of intracellular damage (*8, 11, 12*), it can be regarded as akin to a soma-germline delineation (*13*), effectively creating a separation between an aging mother cell and a rejuvenated daughter with each division. Models on asymmetry in single-cell organisms underpin these findings by showing that asymmetrical segregation of damage is likely to evolve as a strategy to increase fitness in the face of high levels of cellular damage (*14*–*16*). While these models provide a mechanistic understanding of cellular aging, they rely on few empirical accounts on the nature of this damage, its partitioning dynamics, and physiological impacts.

The accumulation of misfolded proteins is widely regarded as an age-associated source of intracellular damage. Sources of proteotoxic stress can cause proteins to change from their native conformation to a misfolded one, rendering them inactive or even toxic to the cell. The misfolding of a protein can expose its hydrophobic portions, leading it to bind to other proteins and thereby interfere with their functioning (*17*). This disruption of proteostasis can lead to the formation of insoluble protein aggregates within a cell. In mammals, the increased accumulation of misfolded proteins and decreased repair capacity is considered an aging hallmark (*1*), often leading to the development of aging-related diseases such as Parkinson’s and Alzheimer’s (*18*). In a striking parallel with eukaryotic processes, *E. coli* aging has been attributed to protein aggregation. Misfolded proteins, which can appear anywhere within the cell, tend to cluster together. If such aggregates become too large to diffuse through the nucleoid (*12, 19, 20*), they can become stuck at the cell pole as it is successively inherited across generations, thus being preferentially retained by the mother cell (*8, 11*). Since the presence of aggregates hinders diffusion of other intracellular components (*19*), it might also cause the mother cell to inherit fewer ribosomes (*21*) and newly synthesized proteins (*22*). Daughter cells, in contrast, inherit a larger share of “good” components along with smaller damage loads, leading to improved rates of growth and gene expression (*23*).

Proteostasis is essential for cellular functioning, and protein repair is key for healthy cell growth. The protein repair machinery of bacteria deals with damage by degrading or refolding misfolded proteins, thereby preventing aggregation, or by disaggregating already formed aggregates (*24*). As such, protein aggregation mainly occurs when a cell accumulates damage at a faster rate than it can repair (*11*), at which point *E. coli* cells exhibit increasing amounts of polar aggregates. Computational analyses suggest that damage aggregation can be advantageous in face of stress, as it allows for the asymmetric partitioning of intracellular damage between aging mothers and rejuvenating daughters (*25*). If the same damage loads were to be symmetrically partitioned, the preservation of cellular functions would likely require a larger investment into repair activities.

While it is known that protein aggregates form in response to stress, their correlation with aging phenotypes has been questioned by recent studies. The observation of large aggregates in maternal old poles has been attributed to the choice of fluorescent marker in earlier research, which used the small chaperone Inclusion Body Protein A (IbpA) labeled with yellow fluorescent protein (YFP) for damage quantification (*8, 26*). When the reporter was exchanged for a monomeric superfolder green fluorescent protein (msfGFP), heat shock-induced aggregates had no negative impact on fitness (*27*). Similarly, *E. coli* lineages grown in unstressed conditions did not accumulate polar aggregates, despite consistently exhibiting a physiological asymmetry between mothers and daughters (*28*). Once the formation of aggregates is induced, however, they can have a protective effect against further heat shock (*27*) and promote antibiotic persistence (*29*), suggesting that their presence does not always imply a fitness decline.

The observation of an age-related loss of proteostasis in prokaryotes has framed this process as a conserved aging hallmark among cellular organisms (*30*). It is therefore imperative to elucidate the recent conflicting evidence on the correlation between protein aggregation and bacterial aging. To address this, we quantified the accumulation of intracellular damage using DnaK, a heat-shock protein that plays a central role in protein quality control, acting both to refold soluble misfolded proteins (*31*) and to disaggregate protein aggregates (*11, 17*). As a damage marker, most previous studies have focused on IbpA (*8, 12, 27*), which co-localizes with misfolded proteins. DnaK, in contrast, reports not only the localization of damage, but also indicates that it is being actively repaired by the cell in an energy-consuming process (*11, 32*). Quantifying the accumulation of DnaK-labeled damage, we found that protein aggregates accumulate in maternal old poles over generations, even as mother cells reach the state of long-term growth stability that is commonly observed within microfluidic devices (*9, 28, 33*). By itself, the presence of a polar aggregate did not lead to lower elongation rates. Instead, we found that the main determinant of aging is the relative size of the protein aggregate, compared to the total length of a cell. We found that aging also leads to an increase in maternal cell size, which buffers the effect of damage accumulation and allows the cell to maintain stable elongation rates despite harboring large clusters of misfolded proteins. We thus propose that protein aggregation is a hallmark of bacterial aging, driving a progressive functional decline in a size-dependent manner.

## RESULTS

### *E. coli* expressing DnaK-msfGFP shows wild-type levels of asymmetry

To evaluate whether protein misfolding and aggregation act as a driver of bacterial aging, we quantified damage accumulation at the single-cell level. For this, we used *E. coli* cells expressing DnaK-msfGFP as a reporter for both the subcellular localization of damaged components and the ongoing repair activity that is expected to be energy-costly (*11*). We ensured stable conditions by culturing cells in the mother machine microfluidic device (*33*), containing thousands of growth wells that receive a continuous supply of nutrients. The closed end of each well traps the cell inheriting an old pole with each division, which can be regarded as the mother cell. The opposite pole, synthesized during the previous fission, is considered a new pole that is inherited by the daughter cell. While the device traps mother cells for hundreds of generations, daughters are lost into the flow channel after a few divisions. Here, we followed populations over 48h, measuring growth physiology and damage accumulation in mother cells over time, and in daughter cells over their first cell cycle. Overall, our analysis encompasses 487 cell lineages expressing DnaK-msfGFP across a total of 18,756 division events, and 346 wildtype (WT) lineages over 15,247 divisions.

To determine whether the expression of the fluorescent reporter had an influence on cell growth, we compared the reporter strain to wild-type *E. coli* MG1655. Since our work explores the impact of protein aggregates on physiological asymmetry, we first confirmed that both strains produced daughters with faster elongation rates than mother cells (Fig. 1A; Kolmogorow-Smirnov (K-S) test: DnaK: D = 0.246, p < 0.001; WT: D = 0.253, p < 0.001). By measuring growth asymmetry as the ratio of elongation rates (Fig, 1B), we found that the DnaK-msfGFP strain was slightly less asymmetric (0.875 ± 0.192, mean ± SD) than the wild-type (0.858 ± 0.181; K-S test: D = 0.0466, p < 0.001), suggesting that daughter cells had a lower growth advantage over their mothers when expressing the fluorescent marker. We also observed a small growth difference in mean elongation rates of both strains, with DnaK-msfGFP cells exhibiting a 5.6% slower growth (0.016 ± 0.005 min^-1^) than the wild-type (0.017 ± 0.005 min^-1^; K-S test: D = 0.098, p < 0.001).

**Fig. 1.**
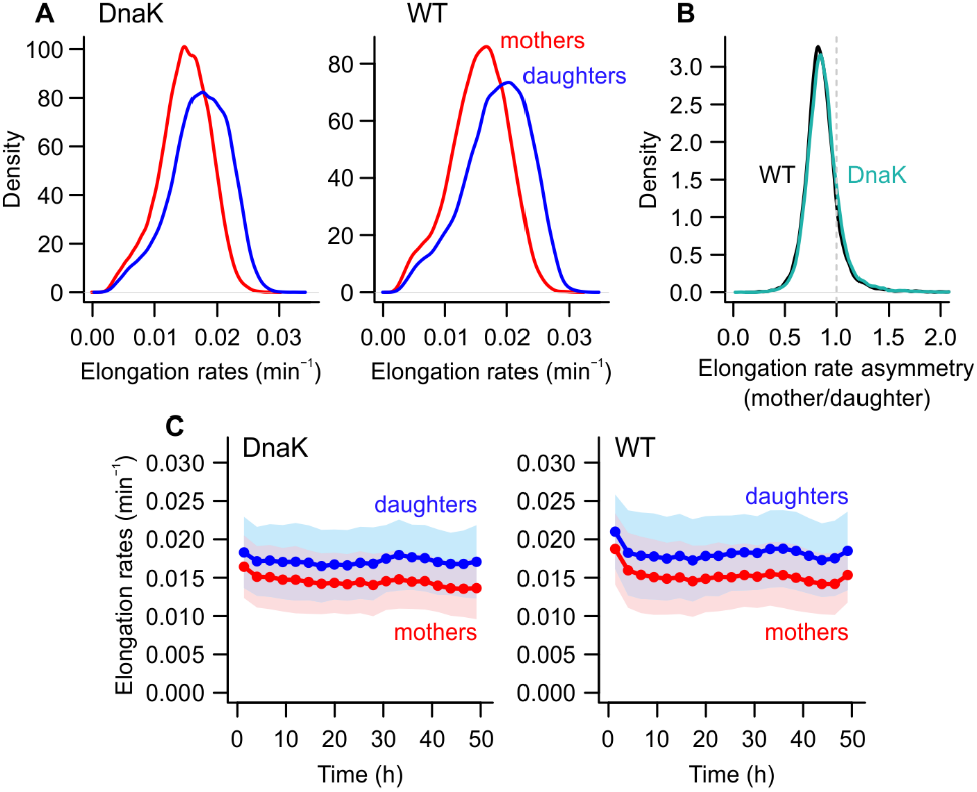
Strains expressing DnaK-msfGFP show asymmetry equivalent to wild-type bacteria. (A) Daughter cells exhibited higher elongation rates for both *E. coli* strains MG1655 *ΔlacY dnaK-msfGFP* (DnaK) and MG1655 *ΔlacY* (WT). (B) Both strains showed similar growth asymmetry, as expressed by mother-daughter elongation rate ratios. (C) In both strains, mother and daughter cells showed stable growth over time, with mothers consistently growing more slowly than daughters. Bins = mean ± SD.

We further evaluated how elongation rate is shaped by time, asymmetry, and expressing the fluorescent fusion through generalized additive models (GAMs; statistical model comparison provided in the Supplementary Text). Whether considering each strain individually or in combination (Tables S1-S3), we found that asymmetry accounts for ∼7.7% of the deviance in the data, indicating that mothers and daughters have significantly distinct intercepts. While accounting for changes in asymmetry across time improved the model fit, this interaction accounted for < 0.2% of the deviance in elongation rates. More importantly, once the models included the effects of time and asymmetry, considering the difference in intercept between the two strains only explained a further 0.4% of the deviance in elongation rates. Since the small effect size of the differences between strains imply a limited biological concern for the context of this work, we conclude that DnaK-msfGFP is a suitable translational reporter for quantifying the correlations between protein aggregation, cell growth, and physiological asymmetry leading to bacterial aging.

### Mean intracellular DnaK concentration increases with age

To determine whether aging correlates with the accumulation of misfolded proteins, whether soluble or aggregated, we measured the mean DnaK-msfGFP fluorescence after each division. We found that the levels of intracellular damage in mother cells increased over time, which was contrasted by no relevant increase in daughter cells (Fig. 2A). We verified this statistically through the comparison of multiple GAMs (Table S4). This analysis showed that asymmetry leads to distinct mean fluorescence levels in mothers and daughters, which varied independently across time. Considering both this mean effect and the interaction with time, asymmetry accounted for 7.3% of the deviance in mean fluorescence. As an individual effect, mean changes across time explained another 7.1% of the deviance in the data. The variance among maternal lineages was largely meaningful (deviance explained = 42.8%) for DnaK-msfGFP levels, suggesting that once a lineage reaches a certain concentration of misfolded proteins, the mother cell is likely to retain this damage over successive divisions.

**Fig. 2.**
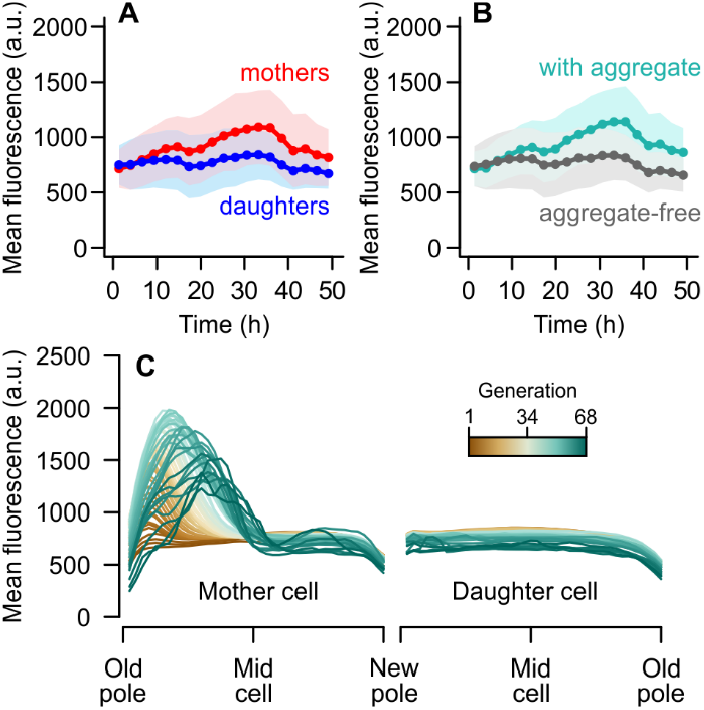
Fluorescence increases over time in mother cells due to aggregates. (A) Mean fluorescence increases over time in mother cells but not in daughters. (B) When the population is divided according to the presence/absence of aggregates, they show a similar increase in mean fluorescence over time as (A), suggesting strong overlap of mother-cell and aggregate-bearing subgroups. Bins = mean ± SD. (C) Mean DnaK-msfGFP fluorescence profiles of mother (left) and daughter cells (right) from old to new pole. Each line represents the average transect for a given generation over a normalized cell length. The mother shows increased GFP levels, indicating protein aggregation, near the old pole.

The asymmetric retention of damage depends on both the formation of protein aggregates and their retention in maternal old poles. To investigate the relationship between asymmetry and protein aggregation, we repeated our analysis while splitting the population according to the presence of clustered damage (Fig. 2B), independently of polar age. The resulting difference between aggregate-bearing and aggregate-free cells largely reflected that of mother-daughter asymmetry (Table S5), explaining a total of 8.15% of the deviance in mean fluorescence. This similarity suggests that the increase in maternal intracellular damage was due to the accumulation of protein aggregates, which are mostly absent among daughter cells.

Next, to determine whether these aggregates were associated with maternal old poles, we quantified the intracellular distribution of DnaK-msfGFP (Fig. 2C). We traced the mean fluorescence profile along the length of each cell immediately after division, and estimated the effect of time and the distance from the old pole on fluorescence levels (Table S6). While the profiles in Fig. 2C depict the average measurements for all cells at a given generation, statistical analyses were performed on individual transects. For mother cells, we found that fluorescence levels changed across the cell axis, with the shape of this profile changing over time (Table S6). Once we accounted for this interaction between the subcellular localization and time, including individual effects of time on mean fluorescence resulted in little improvement to the deviance explained by the models. This indicates that mother cells undergo a localized accumulation of intracellular damage over generations, while showing small changes to the levels of dispersed fluorescence. Daughter cells, on the other hand, had slight changes in mean fluorescence levels over time and across the cell length, but no interaction between these factors.

Taken together, these data indicate that mother cells retain large clusters of damaged proteins over generations, whereas daughters are usually born free of protein aggregates. Beyond a passive accumulation of damage in maternal old poles, this asymmetric inheritance implies an active process: since DnaK acts to refold and disaggregate damaged proteins, consuming ATP in the process, the data in Fig. 2 also suggests an asymmetric allocation of resources where mother cells show a greater investment into repair activities.

### Size of maternal aggregates increases over time

To further investigate the dynamics of protein misfolding and aggregation over generations, we quantified the growth of protein aggregates in maternal lineages. For this, we evaluated fluorescence transects showing the distribution of DnaK-msfGFP throughout the cell’s longitudinal axis after each division. We detected aggregates as fluorescence peaks on each profile (see Methods for details), quantifying their diameter, intracellular position, and mean intensity. As suggested by the average transects in Fig. 2C, we observed increasingly large peaks associated with maternal old poles (Fig. 3A). The aggregates showed an average distance of 0.22 µm from the old pole (Fig. 3B), which represents 7.9% of the mean length of a mother cell at beginning of a cell cycle (2.82 µm). In total, mother cells harbored aggregates on 66.9% of the observed division events. Daughter cells, on the other hand, showed homogeneous levels of intracellular fluorescence (Fig. 2C, right panel), with aggregates being detected only on 3.6% of the quantified transects.

**Fig. 3.**
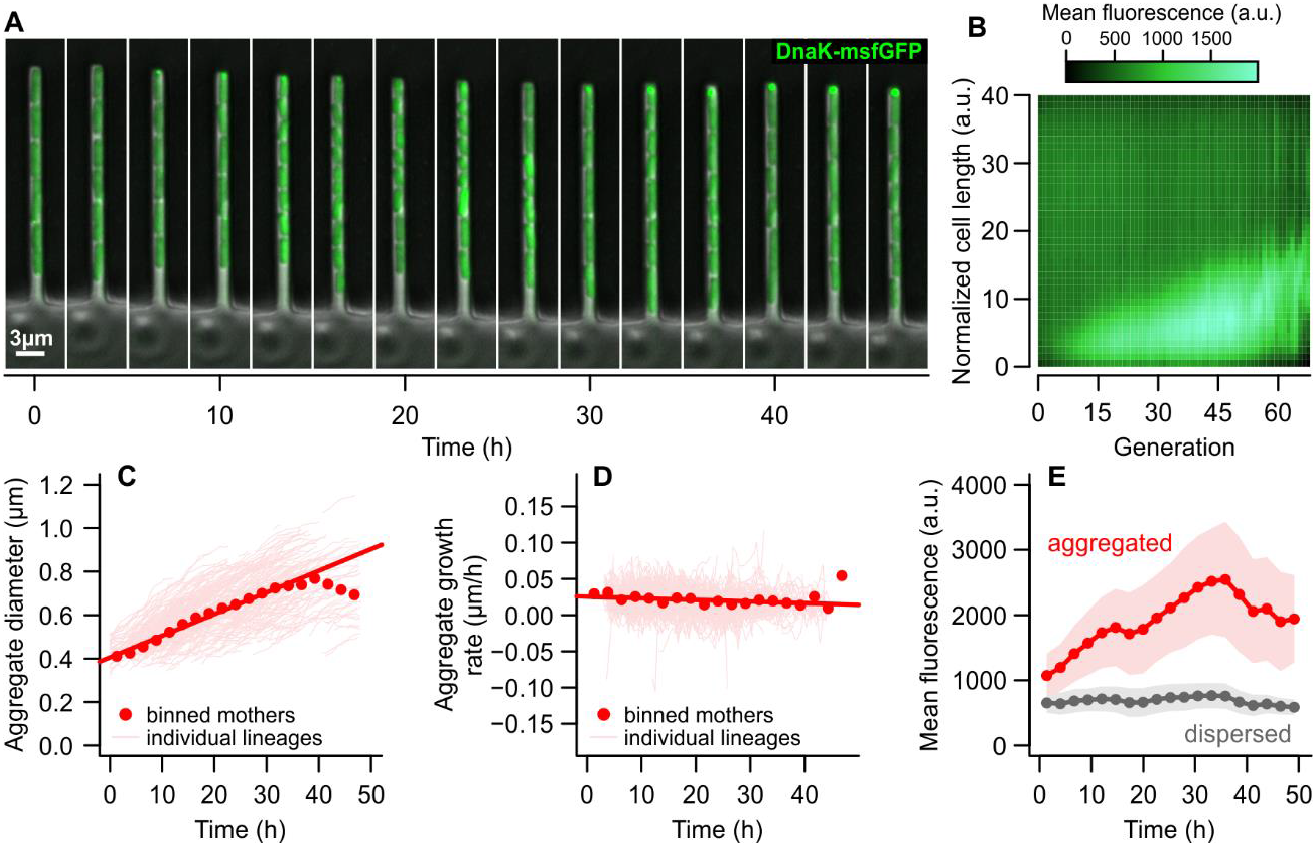
Dynamics of damage accumulation within mother cells. **(A)** Time-lapse images of a cell lineage growing within the mother machine device. The mother cell, at the closed end (top), accumulates DnaK-msfGFP into fluorescent foci, while its daughters (towards the open end of the growth well) exhibit dispersed fluorescence. (B) Heatmap of the mean fluorescence of mother cells across generations, showing that the concentration of DnaK-msfGFP increases in the old poles due to the formation of aggregates. (C) Pooling all aggregate-bearing mothers, we observed that the diameter of protein aggregates increases at a linear rate over time (solid red line, R^2^ = 0.384, p < 0.001). (D) Aggregate diameters changed at a constant rate from one generation to the next, for rates individually estimated for each mother lineage. (E) DnaK-msfGFP concentration in the form of aggregates or dispersed fluorescence in aggregate-bearing mothers. While aggregates increased in size and fluorescence intensity over time, there was little variation in the concentration of misfolded proteins in the rest of the cell. Thus, mother cells accumulate damage at a constant rate as they age, corresponding to the clustering of misfolded proteins into polar aggregates. (C to E) Bins = mean ± SD.

As suggested by the fluorescence profiles in Fig. 3B, maternal aggregates grew larger over time (Fig. 3C and Table S7). Evaluating the growth rate of aggregates across all mother cells, we observed a linear increase in diameters prior to 40h of exponential growth at a rate of 0.01 µm/h (Size = 0.01*Time + 0.4, p < 0.001). Aggregate growth seems to become non-linear after 40h, at which point their diameters are expected to reach 0.8 µm — the maximum width a cell can have within our microfluidic device. To determine whether these growth rates were stable, we estimated the changes in aggregate diameters between consecutive generations for each maternal lineage (Fig. 3D). Although aggregate growth rates fluctuated slightly, any changes over time were negligible (Table S7). By fitting a GAM that evaluated the effect of time and variation among mother cells on aggregate growth, we found that these variables explained 0.09% of the deviance in the data, indicating that aggregates grow in diameter at a stable rate that is subjected to stochastic fluctuations. Interestingly, the linear increase of aggregate diameters with age suggests a buffering effect of aggregation: if we assume that aggregates are spherical, a linear growth in diameter indicates a quadratic change in volume (see Supplementary Text). The quantity of misfolded proteins added to the aggregate would also need to increase quadratically over time, thus implying that a quadratic accumulation of damage only has a linear effect of aggregate diameter growth.

Next, to determine whether mother cells also accumulate non-aggregated misfolded proteins, we quantified the concentration of DnaK in aggregates versus the rest of the cell (Fig. 3E). For this, we determined the intracellular area occupied by fluorescence peaks and the area free of aggregates, classifying the mean fluorescence of the former as “aggregated” and the latter as “dispersed”. We found that, while mother cells accumulated misfolded proteins in the form of aggregates (F = 1310.87, p < 0.001), their levels of dispersed damage showed little change over time (F = 63.72, p < 0.001; Table S8). Together, these results indicate that the accumulation of damaged proteins as mother cells age is strongly associated with their deposition into aggregates, whereas the damage dispersed in the remaining intracellular space stays at baseline levels. We corroborate that the formation of polar aggregates across generations allows for the retention of damage by the mother cell, leading to the asymmetric partitioning of misfolded components upon division.

### Dispersed damage is more harmful than protein aggregates

The increase in aggregate size over time contrasts with the stability of bacterial elongation rates across generations, found in our results (Fig. 1C) as well as in many previous studies (*9, 28, 33*). Whether through a toxic effect of harboring misfolded proteins or through the increased investment of resources into damage repair, it has been widely assumed that protein aggregation should correlate with slower growth (*8, 11*). Past models have proposed that mother cells reach a state of growth stability once the intracellular damage load retained after each division becomes constant (*15, 26*). However, our data suggests that damage levels continue to climb long after mother cells have reached this state of equilibrium (Fig. 1C and Fig. 3C). We must therefore investigate whether the accumulation of misfolded proteins correlates with a physiological decline (*8*) or has no effect on growth (*27*).

Considering the whole population, the presence of a protein aggregate reduced elongation rates, albeit effect sizes were small (Fig. 4A): cells bearing an aggregate grew slightly more slowly (0.0156 ± 0.0039 min^-1^) than aggregate-free cells (0.0162 ± 0.0049 min^-1^; t = -24.45, p < 0.001) (see also Table S9). However, this difference did not factor the inheritance of old or new poles upon division. By repeating this analysis while accounting for the asymmetry between mother and daughter cells, we found that this trend was surprisingly reversed (Fig. 4BC). Aggregate-free cells exhibited slightly slower elongation rates than aggregate-bearing cells in either subpopulation (Table S10; GAM #6: t = 18.12, p < 0.001). Most of the difference in elongation rates could be attributed to mother-daughter asymmetry (t = 50.46, p < 0.001), whereas interactions between time and asymmetry or the presence of aggregates contributing marginally to explain the deviance in elongation rates (Table S10). Despite the small difference between aggregate-bearing and aggregate-free cells, this suggests that the simple presence of an aggregate does not imply slower growth. Once we account for the underlying asymmetry between mothers and daughters, harboring an aggregate might even provide a slight growth advantage.

**Fig. 4.**
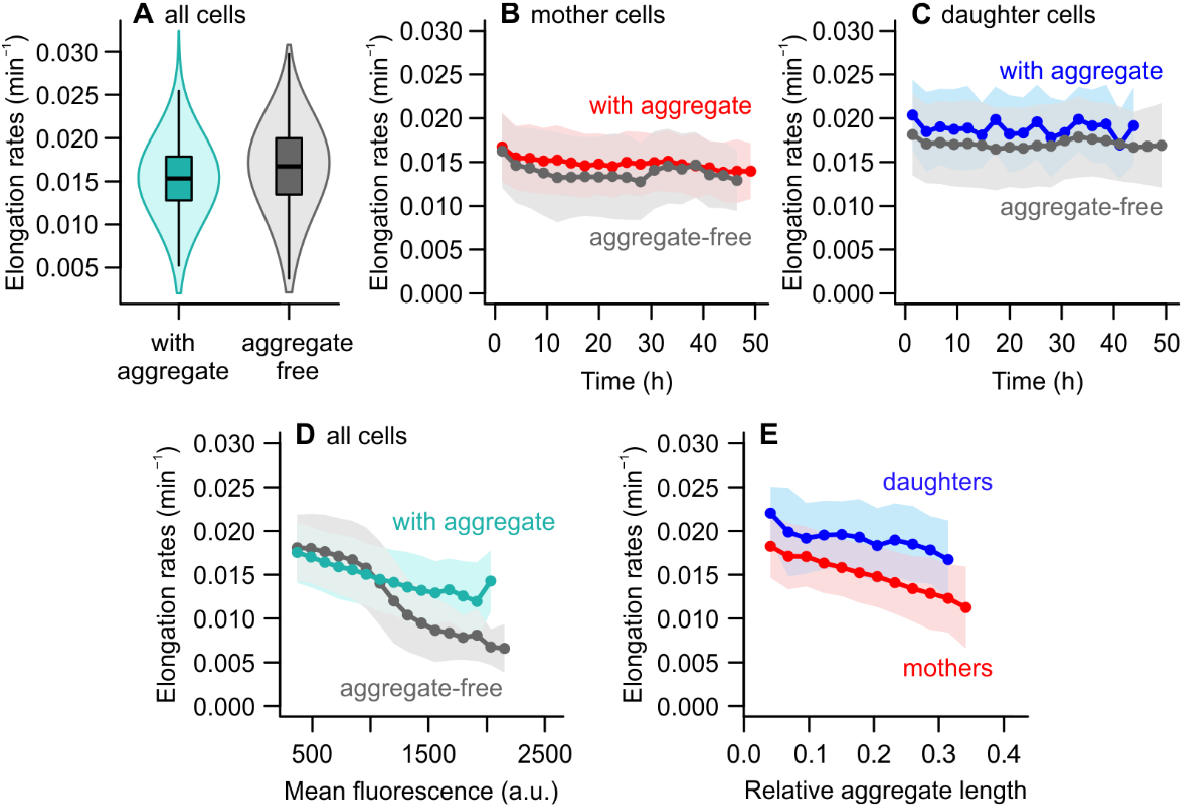
Effect of protein aggregates on cell growth. (A) When considering the population as a whole, elongation rates were negatively impacted by the presence of protein aggregates. (B and C) Splitting the sample into subpopulations of mothers and daughters showed the opposite effect. Within subpopulations, aggregate-bearing cells had consistently greater elongation rates than aggregate-free cells. (D) Intracellular damage loads, as indicated by mean fluorescence, were negatively correlated with elongation rates. High damage concentrations (above 1,000 a.u.) led to a sharp growth decline in cells without aggregates, indicating that dispersed misfolded proteins might be more harmful to the cell at high damage levels. (E) The impact of aggregates on elongation rates depends on the intracellular space occupied by this damage, expressed as the relative size of the aggregate when compared to the length of the cell. Bins = mean ± SD.

This observation appears to stand in contrast with the progressive accumulation of damage among mother cells (Fig. 2C), and the correlation between the inheritance of old poles and slower elongation rates (Fig. 1C). Could it mean that protein aggregation has a protective effect against damage accumulation? To investigate this, we evaluated the mean fluorescence of all cells in the population. Our time-lapse images suggested that, in some cases, growing cells with no discernible aggregates could nonetheless exhibit a high concentration of DnaK-msfGFP. In both mothers and daughters, a fraction of aggregate-free cells showed a mean fluorescence higher than 1,000 a.u., an intensity threshold associated with the formation of a polar aggregate in most other cases (Fig. 2C): fluorescence foci had an average intensity of 1,799.4 ± 768.3 a.u., with their peaks reaching 2,305.3 a.u., whereas aggregate-free cells showed a mean fluorescence of 776.5 ± 250.5 a.u.. By plotting the mean fluorescence of aggregate-free and aggregate-bearing subpopulations against their elongation rates, we observed that a clear distinction emerges for higher DnaK concentrations (Fig. 4D). Although higher fluorescence corresponded to a slower growth in both cases, the effect of intracellular damage levels on elongation rates was modified by whether the cells had aggregates (Table S11), explaining 17.1% of the deviance in elongation rates.

These findings suggest that an increased damage load in the cell may be less disruptive to cellular growth when it is focused or aggregated, rather than dispersed throughout the entire cell — especially when its concentration reaches higher levels. This raises an interesting perspective on bacterial asymmetry: so far, the asymmetric partitioning of damage has been considered advantageous in face of high damage accumulation rates, as it might allow for faster population growth than symmetric partitioning (*16*) and lead to differential stress survival (*15, 25*). However, these past models considered that a given damage load would have the same effect on cellular growth whether dispersed or aggregated in the pole. If damage aggregation decreases the cost of bearing this intracellular load, the very mechanism of asymmetry might provide a fitness advantage over symmetrically dividing cells, which face the increased cost of dispersed damage.

### Cell aging is determined by the relative size of aggregates

Recent studies have shown that ribosomes (*21*) and newly synthesized proteins (*22, 34*) are differentially inherited by the daughter cell, largely due to the exclusion of these components from maternal old poles by protein aggregates (*19*). Thus, the advantage of aggregate-bearing cells could derive from the confinement of damage and repair functions to a smaller area near the cell pole, leaving more free space for processes necessary for healthy cell functioning and growth. In other words, the defining feature affecting *E. coli* cell growth might be the amount of aggregate-free space in the cell. We tested this hypothesis by determining the fraction of the cell occupied by an aggregate (*i*.*e*. relative aggregate length) and determined its effect on elongation rates as a function of cell pole inheritance (Fig. 4E). We found a negative effect of relative aggregate length on the elongation rates of mother and daughter cells (Table S12), which differed slightly in how they varied with increasing aggregate sizes. This suggests that protein aggregates drive a physiological decline in mother cells through the spatial competition with other cellular processes.

### Morphological asymmetry buffers against physiological decline

A puzzling discrepancy still emerges from our results. The diameter of protein aggregates accumulating in maternal old poles increases over time (Fig. 2C), with harmful effects on cellular growth depending on their relative size (Fig. 4E). Nonetheless, elongation rates remain stable across generations. In a bacterium that divides with morphological symmetry, where the size of the mother remains constant, it is not possible to reconcile these observations. If, however, mother cells were to balance out the space that is lost to aggregates by growing longer, this might allow the cell to sustain the normal functioning of growth and protein synthesis.

To test this hypothesis, we evaluated the effects of protein aggregates on the length of mother cells at the start of each cell cycle. We found that cell length increased with replicative age (Fig. 5A), and this effect was modified by the presence of protein aggregates: aggregate-bearing mothers showed a steeper increase in length across generations (F = 426.06, p < 0.001) compared to aggregate-free mothers (F = 61.68, p < 0.001; GAM #1 in Table S13). Because daughter cells do not undergo such changes in length, a morphological asymmetry between mothers and daughters emerges (Fig. 5B). The length asymmetry becomes more pronounced with each division, a pattern that is shaped by the presence of aggregates in mother cells (F = 334.75, p < 0.001; GAM #1 in Table S14). This effect can be attributed to the displacement of the FtsZ fission ring by aggregates (*35*), leading to off-center divisions. In fact, we found that the larger the diameter of the aggregate, the greater the morphological asymmetry (Fig. 5C; GAM: F = 215.4, p < 0.001).

**Fig. 5.**
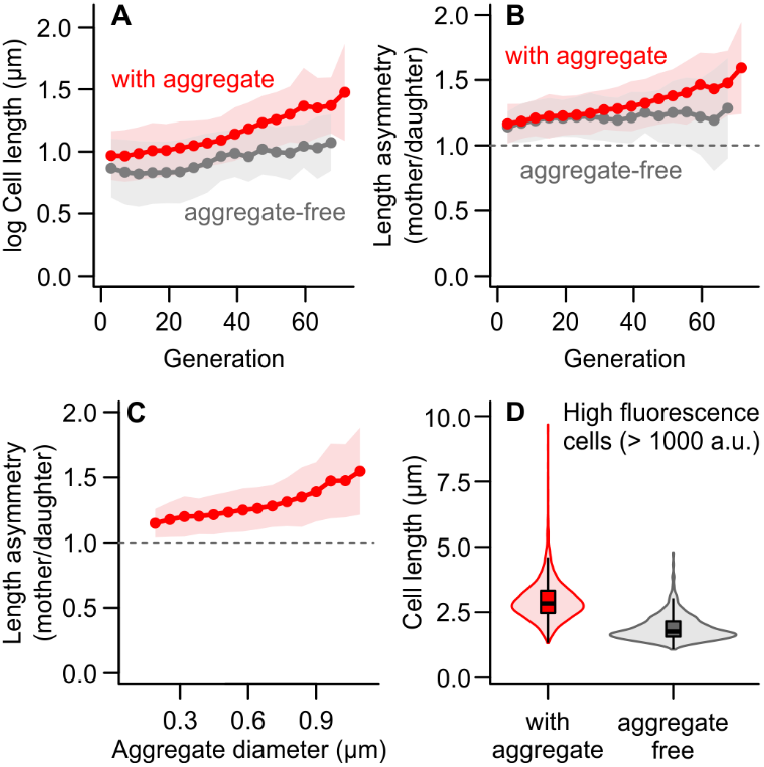
Cell length increases in the presence of protein aggregates, driving length asymmetry. (A) With increasing replicative age, *E. coli* mothers increased in length after division (*i*.*e*., start of the cell cycle). The effect of age on length was greater when an aggregate was present. (B) Since daughter cells do not undergo the same changes in cell length, a morphological asymmetry emerges as mother cells age. This length asymmetry increased with age in aggregate-bearing cells. (C) In divisions where the mother cell carried an aggregate, length asymmetry correlated positively with the size of the aggregate. (A to C) Bins = mean ± SD. (D) Among mother cells with high DnaK concentrations, those bearing aggregates were longer than those containing only dispersed damage. Without the buffering effect of longer lengths, achieved through morphologically asymmetric divisions, the latter suffer a steeper decline in elongation rates for equivalent concentrations of damage (Fig. 4D).

These results support the idea that damage competes for intracellular space with components necessary for cell growth and metabolism. Newly synthesized proteins and ribosomes are known to be excluded from areas occupied by aggregates (*19*) and, consequently, from maternal old poles (*21, 22, 34*). The accumulation of misfolded proteins in the form of aggregates thus hinders cell growth according to the intracellular space occupied by this damage (Fig. 4E). At the same time, however, the formation of aggregates leads to the elongation of the mother cell (Fig. 5A). Because the remainder of the cell has a constant baseline damage level (Fig. 3E), this allows the mother to sustain constant elongation rates (Fig. 1C) even as aggregates increase in size (Fig. 3C). The consequences of lacking this buffering mechanism are best exemplified through a comparison between mother cells harboring high mean fluorescence (Fig. 5D): mothers bearing large concentrations of dispersed damage are significantly smaller than those bearing a similar concentration in the form of aggregates (Table S15), and suffer a stronger fitness decline with the increase in damage loads (Fig. 4D).

Taken together, our results indicate that cell length is affected by the presence and size of protein aggregates accumulated in maternal old poles, resulting in the emergence of morphological asymmetry in aging *E. coli*. Because the physiological impact of damage accumulation depends on the intracellular space limitations, this length increase buffers mother cells against the harmful effects of protein misfolding, allowing elongation rates to remain constant.

## DISCUSSION

Bacteria age through the asymmetry between mother and daughter cells upon division, mirroring the separation between a soma and germline in multicellular organisms. This asymmetry is usually attributed to the uneven segregation of damaged proteins, but the impact of protein aggregation on the bacterial aging phenotype is clouded by conflicting results (*17, 36, 37*). The prevalent assumption was that protein aggregates accumulate in maternal old poles over time, leading to a physiological decline (*8, 11*), but recent findings challenged this perspective by revealing a lack of correlation between aggregates and cell growth (*27*).

We addressed these contradictory findings, showing that mother cells gather misfolded proteins into polar aggregates that grow progressively larger over generations (Fig. 3). Although mother cells were likely to retain this damage upon division, the sole presence of an aggregate could not explain their lower elongation rates. Instead, we found that the main factor driving growth decline was the proportion of the intracellular space occupied by the aggregate (Fig. 4E). One possible explanation for this is the spatial competition between damage and the protein synthesis machinery. The intracellular organization of these components is influenced by the nucleoid (*38*), which holds aggregates in the maternal old pole as they become larger (*12*). Protein aggregates themselves have been shown to hinder the diffusion of other intracellular components (*19*), displacing ribosomes (*21*) and newly synthesized proteins (*22*) towards the new pole. With increasing maternal age, the cell develops an intracellular gradient in gene expression, where the old pole has a lower contribution to the overall metabolism (*23*). Therefore, the relative size of a protein aggregate determines the physiological decline of the mother cell, as it competes with processes that are necessary for cell growth and functioning.

Nonetheless, an unexpected outcome emerged when we considered subpopulations of mother and daughter cells separately. Once the underlying age structure was considered, aggregate-bearing cells showed marginally faster elongation rates than aggregate-free cells (Fig. 4). This effect was more pronounced among cells with a high concentration of intracellular damage, suggesting that polar aggregates might be less detrimental to cellular functioning than similar levels of dispersed damage. This implies that protein aggregation can buffer an aging mother cell against a more severe physiological decline with age. One reason for this buffering effect is that a linear growth in aggregate diameters can only be achieved by a quadratic addition of misfolded proteins. Moreover, the formation of polar aggregates can lead to an off-center positioning of the FtsZ ring (*35*), resulting in the gradual increase in maternal length as the cell accumulates more damage (Fig. 5). The result is the emergence of a morphological asymmetry in *E. coli* division as the mother cell ages, mitigating the spatial competition between damaged proteins and other cellular components — a finding that contrasts with the long-assumed morphological symmetry of these bacteria.

Protein aggregation is a well-observed feature of the aging phenotype, and previous research has shown that it can come with fitness advantages — one being asymmetrical segregation of damage during cell division, ensuring rejuvenated, immortal lineages within the population (*14, 26*). The separation of damaged components from other cellular processes may not only benefit the offspring, but also allow the aging cell to sustain stable growth rates. As mother cells accumulate misfolded proteins in the form of polar-localized aggregates, they compensate for the lost intracellular space by increasing in length. Since this gradual elongation occurs through asymmetric fission, it is justified to assume that these cells continue to divide according to the “adder” model (*39*), growing by a constant length between divisions. This would allow aging cells to maintain a state of growth stability over generations (*9, 28, 33*), showing constant elongation rates despite the continuous accumulation of intracellular damage. For *E. coli*, this could mean that aging may not present as a steady growth decline but rather as a prolonged period of stable growth accompanied by focused damage accumulation, which eventually results in rapid decline and death when damage amounts have reached a lethal level. The existence of such mortality threshold has been suggested by mathematical models (*15, 40*) and even estimated from experimental doubling times (*26*), but its direct quantification remains elusive.

Our findings provide a framework through which protein aggregation operates in bacterial lineages. Increasing damage loads accumulate in mother cells, but their detrimental effects can be buffered by a corresponding increase in length that allows for stable growth. These results, however, do not explain all mechanisms of bacterial aging. An aggregate-free mother cell still grows more slowly than its daughter, and an aggregate-bearing mother has lower elongation rates despite being buffered by a length increase. It is possible that mother cells show decreased growth not only due to harboring more damage, which they can compensate through increased length, but also due to the repair investment required to prevent the growth of protein aggregates from outpacing the lengthening of the cell. In addition, further aging mechanisms not linked to the accumulation of misfolded proteins might contribute to maternal aging, which might not be surprising given the known complexity of the aging process even in simple unicellular organisms. Studies performed on *Caulobacter crescentus*, for example, suggest that bacteria might display other forms of aging even when the segregation of protein aggregates is symmetric (*41*). By demonstrating how growth physiology and protein aggregation dynamics interact along the aging process, we provide insights on the underlying mechanisms that drive senescence and asymmetry at the single-cell level. Besides offering guidance for future studies on the complex and interacting mechanisms of protein misfolding, aggregation, and repair, our findings thus contribute to revealing fundamental mechanisms of aging.

## MATERIALS AND METHODS

### Bacterial strains and growth conditions

*Escherichia coli* strains MG1655 *ΔlacY dnaK-msfGFP* and MG1655 *ΔlacY* (wild-type) were developed at KU Leuven, Department of Microbial and Molecular Systems, for a previous study (*27*). *E. coli* MG1655 *ΔlacY dnaK-msfGFP* expresses a DnaK-msfGFP (monomeric superfolder GFP) fusion protein, which allows for the quantification of aggregate formation. Bacterial stocks were streaked into fresh LB plates, and a single colony was inoculated into 5 ml M9 minimal medium (1x M9 salts, 2 mM MgSO_4_, 0.1 mM CaCl_2_) supplemented with 0.4% glucose and 0.2% casamino acids, where it was grown overnight at 37° C with agitation. Before each experiment, 150 µl of the overnight culture was diluted into 15 ml of fresh medium and grown for 2 h at 37°C. For inoculation into microfluidic devices, the medium was supplemented with 1 μg/ml Propidium Iodide (PI) and 0.075% Tween20 (Sigma-Aldrich). PI facilitates the detection of cell lysis by intercalating with nucleic acids, and Tween20 was used to prevent cell adhesion inside the device. Throughout the microfluidics experiments temperatures were kept constant at 37°C.

### Microfluidic device design and fabrication

The mother machine (*33*) is a microfluidic design that allows for the capture of individual bacterial cells inside narrow growth wells, while being fed fresh culture medium provided by wide flow channels. This allows for the capture of time-lapse images of individual cells growing in exponential phase over several generations. Cells at the closed end of the wells are considered “mothers” because they inherit the old pole in every division. Their new pole siblings, which get washed out of the well into the flow channels over time, are referred to as daughters. Our microfluidic design contained four parallel flow channels (1 cm x 80 μm x 10 μm) with inlet openings on one side and outlets on the other. At these inlets and outlets, metal connectors were connected to Tygon tubing (ND 100-80, Saint-Gobain, Darwin Microfluidics) to feed M9 media into the device and remove cells that had been washed out of the wells. Each flow channel contains 1,000 growth wells (25 x 0.8 x 1.2 μm) that are long enough for mothers not to get washed out too easily, while being short and wide enough for sufficient growth medium to reach mother cells.

We fabricated chips by pouring Sylgard 184 polydimethylsiloxane (PDMS, Dow Corning, USA) into epoxy molds that had been cast from original laser-etched wafers. After pouring, PDMS was degassed to eliminate bubbles introduced during mixing, then cured at 80°C for 60 min. After careful demolding and cropping of the chips, we created inlet and outlet ports at the ends of the flow channels using a 0.5 mm biopsy puncher (Shoney Scientific). Subsequently, chips and ports were washed with ethanol and rinsed with distilled water prior to bonding to 24 x 40 mm coverslips using a plasma cleaner. Bonded chips were immediately placed on a heating plate at 80°C for 2 min, and incubated at 60°C overnight. One hour before cell loading, chips were plasma-activated and flushed with 20% polyethylene glycol (PEG) to aid media flow and prevent cell attachment.

### Cell loading and experimental conditions

Exponential bacterial cultures were centrifuged at 4,000 rpm for 10 min, and the resulting pellet was resuspended into 200 µl of M9 medium. The suspension was injected through the outlet port of each flow channel, followed by the centrifugation of the chip at 1,500 rpm for 8 min to push bacteria into the growth wells. After inspecting the chip under the microscope to ensure that loading was successful, it was connected to inlet tubing running through a peristaltic pump (Ole Dich, Denmark) to generate a constant flow of 200 µl/h. At the outlet ports of the loaded chip, another set of tubing was connected to a waste container to collect the outflow. The chip was then placed in the microscope’s temperature-controlled incubation chamber (Pecon TempController 2000-1, set to 37°C) for the duration of each experiment.

### Microscopy and image acquisition

Time-lapse images were captured with a DS-Qi2 camera connected to an inverted microscope (Nikon Eclipse Ti2) equipped with a motorized stage and Perfect Focus System (PFS). Using a 100x oil objective and the NIS Elements software, once a chip was mounted in the microscope, we selected 12 fields for automated image capture in 2 min intervals. GFP and RFP fluorescence imaging (400 ms exposure and 8% intensity) was set to 10 min intervals to reduce phototoxic stress while providing sufficient data. Two separate replicates were performed for each strain.

### Image processing

After the experiments, images were first adjusted on ImageJ for chromatic shift correction and background subtraction of fluorescence images (rolling ball radius = 12 px). The tool Deep Learning for Time-lapse Analysis (DeLTA) (*42*) was used for automated image processing, which was previously trained on data from our experimental setup. This procedure was performed on the High-Performance Computing center at FU Berlin (*43*). Phase contrast images were used to perform cell segmentation and tracking. The DeLTA output was then converted into R data serialization (RDS) files and visually inspected for quality control. The data for each bacterial lineage was run through a custom R script (version 2023.12.1+402) to correct obvious tracking errors, remove empty or only partially captured wells, and crop wells where severe tracking and segmentation errors occurred. Where necessary, further manual corrections were performed with the same custom R script.

### Quantification of bacterial growth

Elongation rates were used as a measure of cellular growth and fitness, because the rate at which the cell increases in length is crucial for its ability to double its length and divide. To measure elongation rates, each cell was followed from its “birth” until the point where it divided again, and the division interval was determined. Cell length was measured along the cell cycle. Based on these measurements and considering that *E. coli* cell growth is exponential in nature, elongation rates were calculated as the exponential fit of the length increase of each cell over time. As mother and daughter cells originating from the same division were paired in the data, we were also able to determine morphological asymmetry by using length measurements after division to calculate the ratio of mother cell birth length to daughter cell birth length.

### Quantification of dispersed fluorescence and protein aggregates

To determine intracellular damage and protein aggregates, the first fluorescence image of each cell cycle was measured in terms of the brightness of each pixel within the area of the cell. GFP signal intensity was measured as arbitrary units, as fluorescence measurements depend on the microscope and its settings. The same settings were used in all experimental runs, prioritizing resolution at the bright end of the spectrum as we were interested in the areas of the cells with the most fluorescence. To determine the mean intensity of GFP fluorescence and thereby assess the relative amount of DnaK-indicated damage load per cell, an average value over all pixels in the cell was calculated.

From the DeLTA output, we obtained a pixel-by-pixel fluorescence profile along a transect running through the center of each cell, from the old pole to the new pole. In a cell with diffused fluorescence, transect measurements were relatively uniform. However, many cells displayed extreme spikes in fluorescence near the old poles, which could be detected as outliers in the transect data. By locating these outliers using the function findpeaks() in R, areas with a very high density of DnaK-msfGFP, representing aggregates, could be detected and their absolute length, distance to the old pole, intensity peak and mean fluorescence were determined with a custom R script. The ratio of aggregate length to cell length was calculated to gain information about the proportion of the cell which was taken by the aggregate. For visualization of the mean transect measurements over generations, generations with fewer than 10 cells were removed.

### Statistical analysis

All statistical analyses were performed in R (version 2024 4.3.3). Included in the analysis were mother cells as well as their immediate new pole offspring (daughters). These pairs of old and new pole cells born in the same division formed a set of paired data points, each of them observed from its birth until its next division. Any cells that did not complete their division cycle and whose next division was therefore not captured were excluded from the analysis. In the same vein, cells that started their cell cycle before image capture began were also excluded. Mother cell age was determined both in the sense of replicative age (counted in generations, Fig. 2C, Fig. 3B, Fig. 5AB) and chronological age (Fig. 1, Fig. 2AB, Fig. 3C-E, Fig. 4BC), the latter with a resolution of 2 min, as this was the imaging interval. Generation and time count started with the start of imaging.

We used model selection based on AIC (Akaike’s Information Criterium) to differentiate the support of competing models (*44*). We interpret a ΔAIC > 2 as substantially better support. Non-linear patterns were tested using Generalized Additive Models (GAM) (R package mgcv with maximum likelihood estimation). Within each set of models compared, the best-supported model was evaluated for fitting and assumptions using diagnostic plots with the same statistical package. For analyzing mean fluorescence patterns (Fig. 2AB), we used negative binomially distributed error structure with identity link function. For testing interactions between two continuous smoothing parameters (Fig. 2C), we used ‘ti’ (tensor product interaction) rather than the otherwise implicit ‘s’ smoother matrix. For comparison among density distributions Fig. 1AB, we used two-sided two-sample Kolmogorov-Smirnov tests. When evaluating aggregate diameters (Fig. 3A), we were interested in estimating the linear growth and therefore used a linear model. Model assumptions were again evaluated using diagnostic plots. Detailed model estimates are provided in the Supplementary Materials. Data was standardized or normalized as required for visualization, but not for statistical testing. Where plots show binned data, bins represent mean ± standard deviation (SD), excluding bins that contained fewer than 10 data points.

## ACKNOWLEDGEMENTS

We thank L. Stäcker and M. Lajine for experimental support. The authors would like to thank the HPC Service of FUB-IT, Freie Universität Berlin, for computing time. AN was funded by the German Research Foundation (grant 430174701), MT by the Marie Skłodowska-Curie Actions’ European Postdoctoral Fellowship (grant 101069035), UKS by the Heisenberg Programme of the German Research Foundation (grant 430170797), and AMP was funded by the Humboldt Research Fellowship (Alexander von Humboldt Foundation, Germany) and the Rising Star Research Fellowship (FU Berlin, Germany).

