## Supplementary Information for "Protein aggregation drives cell aging in a size-specific manner in *Escherichia coli*"

### SUPPLEMENTARY TEXT

#### Statistical testing using model comparison based on AIC

For the statistical testing among competing models, we use mainly generalized additive models (GAM). AIC (Akaike's Information Criterion) is used to evaluate the relative fit of competing models. We interpret a ΔAIC>2 as better support, with lower AIC indicating better support (Burnham & Anderson 1998), and discuss the potential biological significance of each factor given the deviance explained by it.

#### Comparing DnaK and WT elongation rates across time and cell type (Fig. 1C)

For each strain, we compared the elongation rates of mother and daughter cells (Celltype) across time. Time was used as the smoothing term for the GAM. Fitted interactions are noted as ":", additive terms are noted as "+". The best supported model is in **bold** font. Model assumptions of the best fitting model were assessed via diagnostic plots using the function gam.check() of package "mgcv" in R.

First, we provide two tables that explore elongation rates within the DnaK (Table S1) or the WT strain (Table S2), followed by additional comparisons of both strains combined (Table S3).

Table S1. Effect of time and cell type (mother x daughter) on elongation rates of cells expressing DnaK-msfGFP (Fig. 1C, left panel).

| M.# | Model parameters for GAM | Dev. | DF | AIC | ΔAIC |
|---|---|---|---|---|---|
| 1 | Elongation ~ Time + Lineage* | 14.9% | 450.16 | -286143 | 3233 |
| 2 | Elongation ~ Time + Celltype:Time + Lineage* | 15.1% | 458.37 | -286203 | 3173 |
| 3 | Elongation ~ Celltype + Time + Lineage* | 22.2% | 455.14 | -289309 | 67 |
| **4** | **Elongation ~ Celltype + Time + Celltype:Time + Lineage*** | **22.4%** | **463.42** | **-289376** | **0** |

* as random effects
Dev. = deviance explained

The model comparison for the DnaK strain illustrates the main factors shaping elongation rates in a cell population. While the passage of time alone explained some of the deviance in the data (model 1), accounting for the asymmetry between mothers and daughters increases the deviance explained by 7.28% (contrasting models 1-3 or 2-4). This indicates that elongation rates are best described by different intercepts, which is consistent with the physiological differences expected to emerge from divisional asymmetry. On the other hand, introducing an interaction term between time and Celltype, which would suggest that mothers and daughters vary differently across time, resulted in but a small improvement of the models (0.18% increase in deviance explained, comparing models 1-2 or 3-4). Therefore, while asymmetry itself is an important factor, we may consider changes to this asymmetry across time as negligible.

Elongation rates are also largely shaped by stochastic differences among cell lineages. By considering lineage as a random effect, we comprise each mother cell and its offspring as possibly having a distinct intercept, which increases the deviance explained by 13.3%. This stochastic effect is generally important for the phenotypic heterogeneity of a population, even though it is not the focus of the current work.

In summary, the best-supported model indicated a significant effect of asymmetry on elongation rates ($t = 57.17$, $p < 0.001$, dev. = 7.28%), which vary smoothly across time ($F = 87.04$, $p < 0.001$, dev. = 1.62%) and with stochastic fluctuations among lineages ($F = 11.40$, $p < 0.001$, dev. = 13.3%).

Table S2. Effect of time and cell type (mother x daughter) on elongation rates of WT cells (Fig. 1C right panel).

| M. # | Model parameters for GAM | Dev. | DF | AIC | ΔAIC |
|---|---|---|---|---|---|
| 1 | Elongation ~ Time + Lineage* | 17.0% | 317.70 | -229264 | 2831 |
| 2 | Elongation ~ Time + Celltype:Time + Lineage* | 17.1% | 326.49 | -229293 | 2802 |
| 3 | Elongation ~ Celltype + Time + Lineage* | 24.6% | 321.79 | -232061 | 34 |
| **4** | **Elongation ~ Celltype + Time + Celltype:Time + Lineage*** | **24.7%** | **330.61** | **-232095** | **0** |

* as random effects
Dev. = deviance explained

The model comparison for wild-type cells mirrors qualitative findings for the DnaK strain, illustrating that elongation rates differ between mothers and daughters (models 2 to 4 are better supported than model 1). Accounting for asymmetry led to a 7.6% increase in deviance explained, indicating that mothers and daughters have distinct intercepts. Again, the interaction between Celltype and Time provided only a small improvement (deviance explained = 0.1%).

Table S3. Effect of time, strain (DnaK x WT) and cell type (mother x daughter) on elongation rates.

| M.# | Model parameters for GAM | Dev. | DF | AIC | ΔAIC |
|---|---|---|---|---|---|
| 1 | Elongation ~ Time | 2.81% | 10.90 | -506540 | 6085 |
| 2 | Elongation ~ Celltype + Time | 10.15% | 11.91 | -511603 | 1022 |
| 3 | Elongation ~ Celltype + Time + Celltype:Time | 10.32% | 20.65 | -511709 | 916 |
| 4 | Elongation ~ Celltype + Time + Strain:Time | 10.56% | 20.54 | -511883 | 742 |
| 5 | Elongation ~ Celltype + Strain + Time | 11.04% | 12.91 | -512245 | 380 |
| 6 | Elongation ~ Celltype + Strain + Time + Celltype:Time | 11.21% | 21.66 | -512352 | 273 |
| 7 | Elongation ~ Celltype + Strain + Time + Strain:Time | 11.44% | 21.56 | -512517 | 108 |
| 8 | Elongation ~ Celltype + Time + Strain:Celltype:Time | 10.73% | 37.24 | -511974 | 651 |
| 9 | Elongation ~ Celltype + Time + Strain:Time + Celltype:Time | 10.73% | 28.67 | -511991 | 634 |
| **10** | **Elongation ~ Celltype + Strain + Time + Strain:Time + Celltype:Time** | **11.61%** | **29.75** | **-512625** | **0** |
| 11 | Elongation ~ Celltype + Strain + Time + Strain:Celltype:Time | 11.61% | 38.31 | -512608 | 17 |

Dev. = deviance explained

Combining both strains, we again observed that elongation rates were asymmetric and varied across time. A distinct intercept for mothers and daughters accounted for 7.7% of the deviance explained, largely improving the models (AIC of models 2-10 vs model 1). Including an interaction term between Celltype and time resulted in a small improvement (< 0.2% of the deviance explained), which is consistent with the results observed for each strain independently.

The protein fusion had some influence on elongation rates, as all models that included the strain (models 4-11) had better support than the models that did not differentiate between strains (models 1-3). This improvement was more pronounced when considering a distinct intercept for each strain, whereas accounting for distinct fluctuations across time had a much smaller effect. This is consistent with the slight difference in mean elongation rates observed in Fig. 1, but the model comparison in Table S3 demonstrates that such difference would hardly hold biological significance (deviance explained = 0.89%, comparing models 2&5 or 4&7).

Including the three-way interaction did not improve the model (model 11), suggesting that the elongation rates of mothers and daughters do not vary differently for each strain over time. The best-supported model (model 10) included the two two-way interactions, but this increase in complexity contributed little to the deviance explained: Celltype:Time explained 0.4% of the deviance, and

Strain:Time 0.17%. Therefore, while our large sample size allows for considering multivariate interactions as significant factors acting on elongation rates, there is little gain from this increase in model complexity.

In summary, whereas model 10 is the best-supported model in terms of capturing the small variances in our dataset, model 5 (without interactions) holds the necessary complexity to provide a biologically meaningful interpretation of the data. It indicates that divisional asymmetry is an important factor shaping elongation rates ($t = 72.94$, $p < 0.001$), which show some variance across time ($F = 230.4$, $p < 0.001$) and marginal differences between the DnaK and WT strains ($t = 25.45$, $p < 0.001$).

**Evaluation of DnaK fluorescence signal in mother and daughter cells (Fig. 2)**

In the following analyses, we compared GAMs to determine how the mean concentration of DnaK-msfGFP, as a proxy for accumulated damage, is influenced by time, asymmetry, and the variance between cell lineages. We performed these analyses for separate subpopulations of mother and daughter cells (Table S4), and for cells pooled according to the presence/absence of protein aggregates (Table S5). Finally, we investigate the intracellular distribution of damage observed in Fig. 2C, considering the subcellular localization of DnaK along the cell axis (Table S6).

Table S4. Effect of time and cell type (mother x daughter) on mean fluorescence (Fig. 2A)

| M.# | Model parameters for GAM[a] | Dev. | DF | AIC | ΔAIC |
|---|---|---|---|---|---|
| 1 | Fluorescence ~ Celltype + Time + Lineage* | 53.65% | 488.65 | 475471 | 2652 |
| 2 | Fluorescence ~ Time + Celltype:Time + Lineage* | 51.77% | 496.06 | 476922 | 4102 |
| 3 | Fluorescence ~ Time + Lineage* | 49.61% | 486.84 | 478498 | 5678 |
| **4** | **Fluorescence ~ Celltype + Time + Celltype:Time + Lineage*** | **56.95%** | **498.20** | **472820** | **0** |

[a] fitted with negative binomial error structure
* as random effects
Dev. = deviance explained

The model comparison for fluorescence signals illustrates that the DnaK-msfGFP concentration differed among mother and daughter cells, as models that account for asymmetry (models 1, 2 and 4) are better supported. Contrary to elongation rates, mean fluorescence shows a stronger pattern of changes in asymmetry across time: the interaction between Celltype and time accounts for up to 3.3% of the deviance in the data (model 4 vs model 1). If we consider that asymmetry determines a distinct intercept for mothers and daughters, and that each of these subpopulations shows a distinct variation over time, the combined effect of asymmetry determines 7.3% of the deviance in mean fluorescence.

Considering cell lineage as a random effect leads to a large improvement to the model, as it explains an additional 42.8% of the deviance in mean fluorescence. As we will discuss further ahead, this is largely due to the accumulation of misfolded proteins into aggregates: once they become too large to freely diffuse throughout the cell, they tend to remain at the old cell pole across generations. In other words, once a lineage has acquired a large aggregate, it is very likely that the mother cell will retain this aggregate upon each division.

Table S5. Effect of time and aggregate presence on mean DnaK-msfGFP fluorescence (Fig. 2B)

| M.# | Model parameters for GAM[a] | Dev. | DF | AIC | ΔAIC |
|---|---|---|---|---|---|
| 1 | Fluorescence ~ Aggregate + Time + Lineage* | 58.05% | 490.08 | 459264 | 3029 |
| 3 | Fluorescence ~ Time + Aggregate:Time + Lineage* | 53.33% | 490.12 | 463053 | 6819 |
| 4 | Fluorescence ~ Time + Lineage* | 49.61% | 486.84 | 478498 | 22264 |
| **5** | **Fluorescence ~ Aggregate + Time + Aggregate:Time + Lineage*** | **61.52%** | **499.50** | **456234** | **0** |

[a] fitted with negative binomial error structure
* as random effects
Dev. = deviance explained

By replacing asymmetry with an assessment of whether any given cell had a protein aggregate, we found similar results as described in Table S4. Not only the presence of aggregates influenced the mean fluorescence of the cell, as the change in DnaK-msfGFP concentration over time was shaped by whether misfolded proteins were aggregated or dispersed. In total, protein aggregation accounted for 8.15% of the deviance in mean fluorescence. The large similarity of these results with those shown in Table S4 suggest that the presence of aggregates is largely correlated with the inheritance of either new or old poles, closely corresponding to the effect of asymmetry on DnaK concentrations.

Table S6. Intracellular fluorescence dynamics, given as the effect of the distance from the old pole (distance) over generations on DnaK-msfGFP log fluorescence (Fig. 2C)

| M.# | Model parameters for GAM | Dev. | DF | AIC | ΔAIC |
|---|---|---|---|---|---|
| **Mother Cells** | | | | | |
| **1** | **Fluorescence ~ Distance + Generation + Distance:Generation + Lineage*** | **35.58%** | **515.74** | **735461** | **0** |
| 2 | Fluorescence ~ Distance + Generation + Lineage* | 32.81% | 504.54 | 766988 | 31527 |
| 3 | Fluorescence ~ Distance:Generation + Lineage* | 25.59% | 502.47 | 843583 | 108122 |
| 4 | Fluorescence ~ Distance + Distance:Generation + Lineage* | 35.26% | 511.66 | 739115 | 3654 |
| 5 | Fluorescence ~ Generation + Distance:Generation + Lineage* | 28.02% | 511.37 | 818645 | 83185 |
| 6 | Fluorescence ~ Distance + Lineage* | 30.39% | 495.63 | 793531 | 58071 |
| 7 | Fluorescence ~ Generation + Lineage* | 23.02% | 495.33 | 869023 | 133562 |
| **Daughter Cells** | | | | | |
| 1 | Fluorescence ~ Distance + Generation + Distance:Generation + Lineage* | 61.20% | 516.63 | -241941 | 8675 |
| **2** | **Fluorescence ~ Distance + Generation + Lineage*** | **61.67%** | **505.63** | **-250616** | **0** |
| 3 | Fluorescence ~ Distance:Generation + Lineage* | 50.72% | 503.45 | -72029 | 178587 |
| 4 | Fluorescence ~ Distance + Distance:Generation + Lineage* | 60.27% | 510.48 | -225223 | 25393 |
| 5 | Fluorescence ~ Generation + Distance:Generation + Lineage* | 52.17% | 512.12 | -93179 | 157437 |
| 6 | Fluorescence ~ Distance + Lineage* | 60.22% | 496.63 | -224348 | 26268 |
| 7 | Fluorescence ~ Generation + Lineage* | 52.13% | 496.55 | -92741 | 157875 |

* as random effects
Dev. = deviance explained

The fluorescence transects shown in Fig. 2C comprise mean intensity measurements across each pixel of mothers and daughters immediately after every division, which renders a challenging complexity to these models. We therefore do not report on all model comparisons and only present those that are correctly estimated. For reasons of parameter estimation limitation, we refer to specialized literature (Wood et al. 2015, Hauenstein et al 2018).

The occurrence and growth of protein aggregates over generations is best expressed through separate models for mothers and daughters. For mother cells, the best-supported model included the individual effects of distance from the old pole and generation, as well as their interaction. In other words, fluorescence levels varied intracellularly along the cell length, with the shape of these transects changing over time. It is worth noting that, in terms of deviance explained, there was little gain from including the individual effect of Generation (comparing model 1 to model 4): the complete model only explained 0.32% more of the deviance in fluorescence levels. This suggests that the effect of time is better represented by its interaction with the subcellular localization of fluorescence foci, as protein aggregates develop over time in maternal old poles.

For daughter cells, in contrast, the best-supported model did not include an interaction between intracellular localization and time. This illustrates that daughters do not accumulate aggregates in a specific part of the cell, showing instead a slight change in fluorescence over time (mean decrease) and a stable variation across the cell length (corresponding to lower fluorescence levels in the old pole area). Combined, the models presented in Table S6 indicate that protein aggregates accumulate in maternal old poles over time, and are largely absent in daughters cells.

#### Growth of protein aggregates over time (Fig. 3)

Table S7. Linear models of protein aggregate diameter (Fig. 3A) and its growth rate (Fig. 3B) over time, combining all mother cells.

| Model # | Model parameters for linear model | DF | AIC | ΔAIC |
|---|---|---|---|---|
| **Aggregate diameter (µm)** | | | | |
| 1 | Aggregate diameter ~ Time | 3 | -13402 | 0 |
| 2 | Aggregate diameter ~ 1 | 2 | -7917 | 5485 |
| **Diameter growth rate (µm/h)** | | | | |
| 3 | Aggregate growth ~ Time | 3 | -13336.43 | 0 |
| 4 | Aggregate growth ~ 1 | 2 | -13334.19 | 2.24 |

Comparing a linear model that includes the size of protein aggregates over time (model 1) to one that only holds an intercept (model 2) shows that aggregates grow over time in a linear way (diameter = 0.00995*Time + 0.4056, $R^2 = 0.384$, $p < 0.001$). In order to estimate more accurately the growth rate of these aggregates, we right-censored data beyond 40h, as aggregate growth seemed to become non-linear in such very old cells. Qualitatively, the interpretation does not change. The table above presents the findings for the complete, i.e. non-right-censored dataset.

Regarding the linear growth rates of maternal aggregates, the model comparison using ΔAICs indicated a negligible effect of Time (growth rate = -0.00023*Time + 0.0254, $p = 0.04$), indicating that mother cells experience accumulate damage at a stable rate over generations. To further evaluate this trend, we modelled aggregate growth rates through a GAM where Growth rate ~ Time + Lineage, with the latter consisting of a random effect. This model explained only 0.09% of the deviance in the data, indicating no effect of Lineage on damage accumulation rates ($F = 0$, $p = 1$) and a small effect of time ($F = 3.05$, $p = 0.03$). Thus, although aggregate growth rates may vary among lineages and over time, this pattern appears to be stochastic.

Table S8. Effect of damage aggregation (Source: aggregated or dispersed) of DnaK-msfGFP and Time on mean fluorescence levels of aggregate-bearing mother cells (Fig. 3E).

| M.# | Model parameters for GAM[a] | Dev. | DF | AIC | ΔAIC |
|---|---|---|---|---|---|
| **1** | **Fluorescence ~ Source + Time + Time:Source + Lineage*** | **81.7%** | **438.6** | **340099** | **0** |
| 2 | Fluorescence ~ Source + Time + Lineage* | 75.4% | 419.4 | 347305 | 7207 |
| 3 | Fluorescence ~ Time + Lineage* | 20.5% | 385.8 | 376285 | 36186 |

[a] fitted with negative binomial error structure
* as random effects
Dev. = deviance explained

Considering all aggregate-bearing mother cells, we evaluated whether the increase in mean fluorescence over time (Fig. 2A) was solely due to the accumulation of damage in the form of aggregates, or whether there was also an increase in dispersed DnaK-msfGFP signal. The best-supported model (model 1) indicates not only a difference in the intercept, corresponding to the greater fluorescence of aggregated proteins, but also an interaction between the fluorescence Source and Time, suggesting that aggregated and dispersed damage accumulate at distinct rates over time.

#### Effect of protein aggregates on cell elongation rates (Fig. 4)

Table S9. Linear models estimating the effect of aggregate presence on cell elongation rates (Fig. 4A)

| Model # | Model parameters for linear model | DF | AIC | ΔAIC |
|---|---|---|---|---|
| **1** | **Elongation ~ Aggregate presence** | **3** | **-276699.2** | **0** |
| 2 | Elongation ~ 1 | 2 | -276108.2 | 591 |

Comparing a model that differentiates elongation rates between aggregate-bearing and aggregate-free (model 1) to one that only holds an intercept (model 2) suggests that aggregates lower cell elongation rates ($t = -24.45$, $p < 0.001$). Note, this initial simple model does not differentiate among mother and daughter cells (see below).

Table S10. Effect of aggregate presence (Aggregate) on Elongation rates over Time, considering the influence of other potential sources of asymmetry (Celltype: mother vs daughter) (Fig. 4BC).

| M. # | Model parameters for GAM | Dev. | DF | AIC | ΔAIC |
|---|---|---|---|---|---|
| 1 | Elongation ~ Aggregate + Time + Aggregate:Time | 3.2% | 19.8 | -277187 | 2405 |
| **2** | **Elongation ~ Aggregate + Celltype + Time + Aggregate:Time + Celltype:Time + Aggregate:Celltype:Time** | **9.7%** | **32.1** | **-279592** | **0** |
| **3** | **Elongation ~ Aggregate + Celltype + Time + Aggregate:Time + Celltype:Time** | **9.6%** | **20.7** | **-279587** | **5** |
| 4 | Elongation ~ Celltype + Time + Celltype:Time | 8.8% | 19.8 | -279260 | 332 |
| 5 | Elongation ~ Time | 1.5% | 10.7 | -276624 | 2968 |
| **6** | **Elongation ~ Aggregate + Celltype + Time + Celltype:Time** | **9.6%** | **20.8** | **-279585** | **7** |
| 7 | Elongation ~ Aggregate + Celltype + Time + Aggregate:Time | 9.5% | 20.5 | -279533 | 59 |
| 8 | Elongation ~ Aggregate + Celltype + Time | 9.4% | 12.7 | -279503 | 89 |

GAM comparisons indicate that the best-supported models require a differentiation between mother and daughter cells when considering the effect of aggregates on elongation rates. While model 2 was the best-supported, models 3 and 6, which did not account for the three-way interaction, showed nearly equal support, with the interaction term explaining only 0.1% of the deviance in elongation rates. This means that, while accounting for asymmetry is essential, there is no evidence that mothers and daughters respond differently to the presence of aggregates, regarding the variation of their elongation rates over time. A much simpler model, such as model 6, therefore suffices to express the patterns in our data.

Contrasting with the conclusions from Fig. 4A, which suggested that aggregate-bearing cells grow slower, the models in Table S10 indicate that this effect should be attributed to the underlying asymmetry between mothers and daughters, rather than the presence of aggregates. For instance, comparing model 6 to model 4, which does not account for the effect of aggregates, shows that the deviance explained by this effect is of only 0.8%. The difference in elongation rates produced by asymmetry is much more important than the difference between aggregate-bearing and aggregate-free cells (model 1 vs. model 7). Combined daughter cells grow faster than mother cells, and they differ slightly in how elongation rates vary over time, with the Celltype:Time interaction only accounting for 0.2% of the deviance explained. These results support the unexpected trend found in Fig. 4B, indicating that aggregate-bearing cells grow slightly than aggregate-free ones, when one accounts for the effect of asymmetry.

Table S11. Effect of mean DnaK-msfGFP fluorescence (Fluo) on elongation rates, considering the presence or absence of aggregates (Fig. 4D).

| Model # | Model parameters for GAM | Dev. | DF | AIC | ΔAIC |
|---|---|---|---|---|---|
| **1** | **Elongation ~ Aggregate + Fluo + Aggregate:Fluo** | **17.1%** | **15.0** | **-282569** | **0** |
| 2 | Elongation ~ Aggregate + Fluo | 14.7% | 11.1 | -281573 | 996 |
| 3 | Elongation ~ Fluo | 14.5% | 10.1 | -281523 | 1046 |

Elongation rates drop differently with increasing mean fluorescence among aggregate-holding and aggregate-free cells. This finding is indicated by the better support of the model that accounts for the Aggregate:Fluo interaction (model 1) compared to model 2, that only includes additive effects of aggregate presence and mean fluorescence. Not accounting for aggregates (model 3) yielded the least support. Combined, the increase in mean fluorescence along with its distribution, whether dispersed or aggregated, has the strongest impact on elongation rates. This supports the conclusion that aggregate-free cells with high fluorescence levels, *i.e.* cells with a high concentration of dispersed misfolded proteins, pay a high growth cost compared to cells where high fluorescence is detected, but in the form of polar aggregates.

Table S12. Effect of the relative cell length occupied by an aggregate (RelAggLen) on the elongation rates of mother and daughter cells (Celltype) (Fig. 4E).

| M.# | Model parameters for GAM | Dev. | DF | AIC | ΔAIC |
|---|---|---|---|---|---|
| **1** | **Elongation ~ Celltype + RelAggLen + Celltype:RelAggLen** | **13.9%** | **5.0** | **-104315** | **0** |
| 2 | Elongation ~ Celltype + RelAggLen | 13.7% | 6.9 | -104277 | 38 |
| 3 | Elongation ~ RelAggLen | 8.5% | 6.5 | -103556 | 759 |

The relative aggregate length describes the fraction of the intracellular space that is occupied by an aggregate. Elongation rates are lowered differently in mother and daughter cells depending on the fraction of space the aggregate occupies (better support for model 1, with Celltype:RelAggLen interaction), although including this interaction only accounted for 0.2% of the deviance explained by the model. Combined, this shows that elongation rates correlate with the amount of intracellular space that is occupied by the aggregate.

#### Effects of aggregates on cell length and mother-daughter length asymmetry

Table S13. Changes on cell length (log) over time, as a function of aggregate presence (Fig. 5A).

| M.# | Model parameters for GAM | Dev. | DF | AIC | ΔAIC |
|---|---|---|---|---|---|
| **1** | **Length ~ Aggregate + Generation + Aggregate:Generation** | **25.9%** | **14.2** | **-3095** | **0** |
| 2 | Length ~ Aggregate + Generation | 25.3% | 9.8 | -2963 | 132 |
| 3 | Length ~ Generation | 17.0% | 7.3 | -1116 | 1978 |

The models we tested indicate that, while mother cells become larger over generations independently of their damage content (model 3), the presence of aggregates has an important role in shaping the increase in cell size (model 1-2). Aggregate-free mothers are overall smaller than those bearing aggregates, which by itself explains 8.3% of the deviance in the data (model 2 vs model 3). Additionally, the best-supported model also considers an interaction term (model 1), suggesting that the presence of aggregates lead to a sharper increase in cell length over time than is observed in aggregate-free mothers — although the additional deviance explained by this interaction showed a small effect (0.6%).

Table S14. Effect of aggregate presence on morphological asymmetry (Fig. 5B)

| M.# | Model parameters for GAM | Dev. | DF | AIC | ΔAIC |
|---|---|---|---|---|---|
| **1** | **LengthAsym ~ Aggregate + Aggregate:Generation** | **13.7%** | **12.9** | **-13024** | **0** |
| **2** | **LengthAsym ~ Aggregate + Generation + Aggregate:Generation** | **13.8%** | **12.9** | **-13024** | **0.1** |
| 3 | LengthAsym ~ Aggregate + Generation | 12.5% | 8.2 | -12777 | 247.1 |
| 4 | LengthAsym ~ Generation | 11.4% | 7.7 | -12567 | 457.4 |

Mothers and daughters do not divide with morphological symmetry, as the mother tends to be larger than the daughter cells. This difference in length immediately after division is the size asymmetry we explore here. Our model comparison indicates that the presence of aggregates is important for the determination of length asymmetry (models 1-3 vs model 4), which increases over time as the mother cell becomes longer. Models that consider an interaction term, indicating that the presence of aggregates shapes change over generations (models 1-2), receive better support than one that assumes that cells with and without aggregates undergo the same asymmetry changes at distinct mean levels (model 3). Interestingly, once this interaction is considered, the passing of time as an individual factor can be removed from the model with little change to deviance explained (model 1 vs model 2). This model comparison demonstrates that cell morphology become more asymmetric with each division, with the asymmetry increase being more pronounced in aggregate-bearing mothers.

Table S15. Linear models estimating the effect of aggregated versus dispersed misfolded proteins on (log) cell length, considering mother cells with high damage levels (mean fluorescence > 1,000 a.u.).

| Model # | Model parameters for linear model | DF | AIC | ΔAIC |
|---|---|---|---|---|
| **1** | **Length ~ Aggregate presence** | **3** | **18.2** | **0** |
| 2 | Length ~ 1 | 2 | 211.6 | 2193 |

Comparing mother cells bearing large damage concentrations, whether in the form of aggregates or dispersed damage, shows that the accumulation of misfolded proteins has an effect on cell length (model 1: $t = 52.2$, $p < 0.001$). Immediately after division, aggregate-bearing mothers are longer than those containing a similar concentration of dispersed damage.

#### Converting diameter growth of aggregate length to volume growth of aggregates

Taking the growth of aggregates from Fig. 3A prior to 40h reveals a linear increase in the diameter of aggregates. The linear regression model for this increase computes an intercept of 0.4056 µm (can be seen as the initial diameter at the detection threshold) and a slope of 0.0099 µm/h (aggregate growth rate). However, our primary focus is not the growth in diameter but in volume, as aggregates can be assumed as spherical. This volume growth should relate to the number of misfolded proteins that are produced and added to the aggregate. The misfolded protein production rate should express the underlying rate at which mother cells accumulate damage, therefore providing an estimate of their aging rates.

We can estimate the aggregate volume $V = \frac{4}{3}\pi r^3$ with $r$ being the radius of the sphere and $V$ being the volume.

Our estimates from Fig. 3 provide intercept ($a$) and slope ($b$) terms relative to aggregate diameters. As the radius is just half of that, the growth function can be expressed as

$$r(t) = a + bt = 0.2028 + 0.00495t \quad (1)$$

with the time step $t$ expressed in hours.

What we aim at, is the change in volume over time $t$,

$$\frac{dV}{dt} = \frac{dV}{dr}\frac{dr}{dt} \quad (2)$$

by estimating the change in the radius, $r$, over time period $t$,

$$\frac{dr}{dt} = b \quad (3)$$

$$\frac{dV}{dr} = 4\pi r^2 \quad (4)$$

If we insert (3) and (4) in (2) we get,

$$\frac{dV}{dt} = 4\pi r^2 b = 4\pi b(a + bt)^2 \tag{5}$$

$$4\pi b(a + bt)^2 = 4\pi 0.00495(0.2028 + 0.00495t)^2 \tag{6}$$

This illustrates that the volume, and therefore the underlying misfolded protein production rate, increases quadratically.
